# Single mutation makes *Escherichia coli* an insect mutualist

**DOI:** 10.1101/2022.01.26.477692

**Authors:** Ryuichi Koga, Minoru Moriyama, Naoko Onodera-Tanifuji, Yoshiko Ishii, Hiroki Takai, Masaki Mizutani, Kohei Oguchi, Reiko Okura, Shingo Suzuki, Yasuhiro Goto, Tetsuya Hayashi, Masahide Seki, Yutaka Suzuki, Yudai Nishide, Takahiro Hosokawa, Yuichi Wakamoto, Chikara Furusawa, Takema Fukatsu

## Abstract

We report an experimental system in which *Escherichia coli* evolves into an insect mutualist. When the essential gut symbiont of the stinkbug *Plautia stali* was replaced by *E. coli,* a few survivor insects exhibited specific localization and vertical transmission of *E. coli.* Through trans-generational maintenance with *P. stali*, several hyper-mutating *E. coli* lines independently evolved host’s high adult emergence and improved body color. Such “mutualistic” *E. coli* lines exhibited independent mutations disrupting the carbon catabolite repression (CCR) global transcriptional regulator. Each of the mutations reproduced the mutualistic phenotypes when introduced into wild-type *E. coli*, confirming that the single CCR mutations instantly make *E. coli* an insect mutualist. Our discovery uncovers that evolution of elaborate mutualism can proceed more easily and rapidly than conventionally envisaged.

---

Microbial symbioses are among the major evolutionary drivers underpinning the biodiversity (*1,2*). How ordinary free-living microbes have become sophisticated mutualists is an important but unanswered question in understanding the evolution of symbiosis. To address this fundamental issue, experimental evolutionary approaches may provide valuable insights (*3–8*). Here we report a novel experimental system in which the model bacterium *Escherichia coli* evolves into an insect mutualist, thereby demonstrating that evolution of mutualism can proceed very easily and quickly via disruption of a global transcriptional regulator system.

## *E. coli* potentially capable of symbiosis with *P. stali*

Plant-sucking heteropteran bugs generally possess specific symbiotic bacteria in the midgut, which contribute to their growth and survival via provisioning of essential amino acids and/or vitamins (*9,10*). The brown-winged green stinkbug *Plautia stali* (Hemiptera: Pentatomidae) (Fig. 1a) develops a specialized symbiotic organ consisting of numerous crypts in a posterior region of the midgut (Fig. 1b). The crypt cavities are densely populated by a specific bacterial symbiont of the genus *Pantoea* (Fig. 1c, d). The symbiont is essential for growth and survival of the host insect. Normal insects infected with the uncultivable obligatory symbiont, *Pantoea* sp. A (*11,12*), attained over 70% adult emergence rates (Fig. 1e), smeared the symbiont cells onto the eggs upon oviposition (Fig. 1f), and transmitted the symbiont vertically to the offspring via nymphal probing of the eggshell (Fig. 1g). Aposymbiotic insects generated by egg surface sterilization died out with no adult emergence (Fig. 1e). Non-symbiotic bacteria, such as *Bacillus subtilis* and *Burkholderia insecticola,* cannot establish infection and symbiosis with *P. stali* (*11*). Meanwhile, when *E. coli* was inoculated to sterilized newborn nymphs, the insects certainly exhibited retarded growth and high mortality, but a small number of adult insects emerged, attaining 5-10% adult emergence rates (Fig. 1e; fig. S1) (*11*). Such adult insects, which were dwarf in size and dark in color (Fig. 1h), tended to die early, but some insects managed to survive, mate, and produce a small number of eggs. We dissected and inspected these insects, and found that, surprisingly, although the symbiotic organ was atrophied (Fig. 1i), *E. coli* localized to the midgut crypts just like the original symbiont, although the infection patterns were often patchy (Fig. 1j, k; fig. S2). Furthermore, *E. coli* cells were smeared on the eggshell and vertically transmitted to the offspring (Fig. 1l, m), although the transmission rates and the infection titers were unstable in comparison with those of the original symbiont (Fig. 1l). These results suggested that, though incipiently, *E. coli* is capable of localized infection, vertical transmission, and supporting host survival in *P stali.* Considering that *E. coli* belongs to the same Enterobacteriaceae as the original *Pantoea* symbiont, *E. coli* may be able to co-opt the mechanisms for infection and localization of the symbiont to establish the incipient symbiosis (*11*). In this context, it seems relevant that, in the stinkbug family Pentatomidae, the gut symbiotic bacteria have evolved repeatedly from the Enterobacteriaceae through recurrent acquisitions and replacements (*13,14*).

**Fig. 1.**
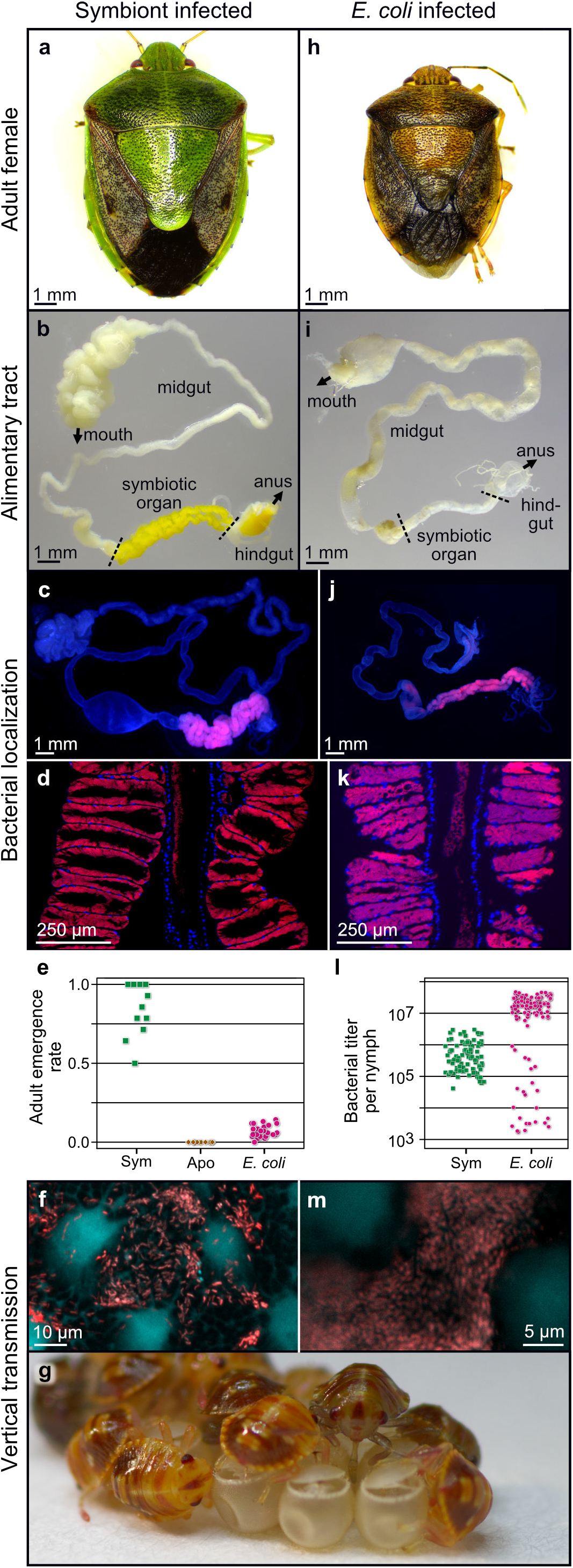
Infection, localization and vertical transmission of *E. coli* in the gut symbiotic system of *P. stali.* (**a**) Normal symbiotic adult female, large in size and green in color. (**b**) Dissected alimentary tract, in which symbiotic organ is well developed and yellow in color. (**c**) Fluorescence in situ hybridization (FISH) localization of symbiont cells to the symbiotic organ. (**d**) Magnified FISH image showing symbiont localization to crypt cavities of the symbiotic organ. (**e**) Adult emergence rates of newborn nymphs inoculated with normal symbiont (Sym, *Pantoea* sp. A), no bacteria (Apo, aposymbiotic), and *E. coli*. (**f**) Symbiont cells smeared on egg surface. (**g**) Newborn nymphs sucking symbiont cells from eggshell. (**h**) *E. coli*-infected adult female, dwarf in size and brown in color. (**i**) Dissected alimentary tract, in which symbiotic organ is atrophied. (**j**) FISH localization of *E. coli* to the symbiotic organ. (**k**) Magnified FISH image visualizing *E. coli* localization to crypt cavities of the symbiotic organ. (**l**) Bacterial titers in symbiont-inoculated and *E. coli*-inoculated nymphs one day after second instar molt in terms of groEL and nptII gene copies per insect, respectively. (**m**) *E. coli* cells smeared on egg surface.

## Experimental evolution using hyper-mutating *E. coli*

This finding prompted us to apply experimental evolutionary approaches to the *P. stali*-*E. coli* relationship. By continuously inoculated to and maintained with *P stali,* would *E. coli* improve the symbiosis-related traits and finally evolve into a symbiont-like entity? Considering the expected difficulty in observing the evolution of elaborate symbiosis in a realistic time frame, we adopted the hyper-mutating *E. coli* strain, ΔmutS, in which the DNA mismatch repair enzyme gene mutS is disrupted and the molecular evolutionary rate is elevated by two orders of magnitude (*15*). The *E. coli* strain of the same genetic background, ΔintS, in which the phage integrase gene is disrupted, was used as control. Two selection schemes, growth selection and color selection, were conducted (fig. S3). In growth selection lines (GmL for hyper-mutating ΔmutS lines; GiL for non-mutating ΔintS lines), the first-emerged adult insect was subjected to dissection of the symbiotic organ for inoculation to the next generation as well as freeze storing. In color selection lines (CmL for ΔmutS lines; CiL for ΔintS lines), the most greenish adult insect was subjected to dissection of the symbiotic organ for inoculation to the next generation as well as freeze storing. Throughout the evolutionary experiments, the host insects were supplied from a mass-reared inbred population of *P. stali*, thereby homogenizing the host genetic background and focusing on the evolutionary changes of the *E. coli* side. Since it takes around 1 month for newborn nymphs of *P. stali* to become adults under the rearing condition, it was expected that, ideally, we would be able to run 12 host generations per year. Actually, however, it took almost for two years because (i) the *E. coli*-inoculated insects generally exhibited high mortality and retarded growth, (ii) for keeping the insects under a good condition, frequent care without overcrowding was essential, which limited the manageable number of insects per evolutionary line ranging from 50 to 100, and (iii) consequently, extended generation time and stochastic extinction of the evolutionary lines frequently occurred, which had to be restarted from the frozen *E. coli* stocks.

## Evolution of mutualistic *E. coli*

We established and maintained 12 CmL color selection lines with 11 CiL control lines, and 7 GmL growth selection lines with 7 CiL control lines (Fig. 2a, b). While the control DintS-infected lines almost constantly exhibited low adult emergence rates, some of the hypermutating ΔmutS-infected lines started to produce more adult insects. Notably, in a color selection line CmL05, the adult emergence rate jumped up at generation 7, and the high emergence rates were maintained thereafter (Fig. 2a). In a growth selection line GmL07, the adult emergence rate improved as early as at generation 2, which was maintained thereafter (Fig. 2b). In CmL05 and GmL07, coincident with the improvement of the adult emergence rate, body color of the adult insects improved from dark to greenish (Fig. 2a-c; fig. S4), and furthermore, the colony morphology of *E. coli* changed from large and flat with rich extracellular matrix to small and convex with little extracellular matrix (Fig. 2c). When the frozen stocks of CmL05 and GmL07 were inoculated to *P. stali,* the improved adult emergence rate, the greenish body color, and the small and convex colony shape were reproducibly observed (Fig. 2d, e; fig. S5). These results indicated that some evolutionary lines of hypermutating *E. coli* have developed mutualistic traits for the host insect and that the phenotypic effects are attributable to genetic changes in the evolutionary *E. coli* lines.

**Fig. 2.**
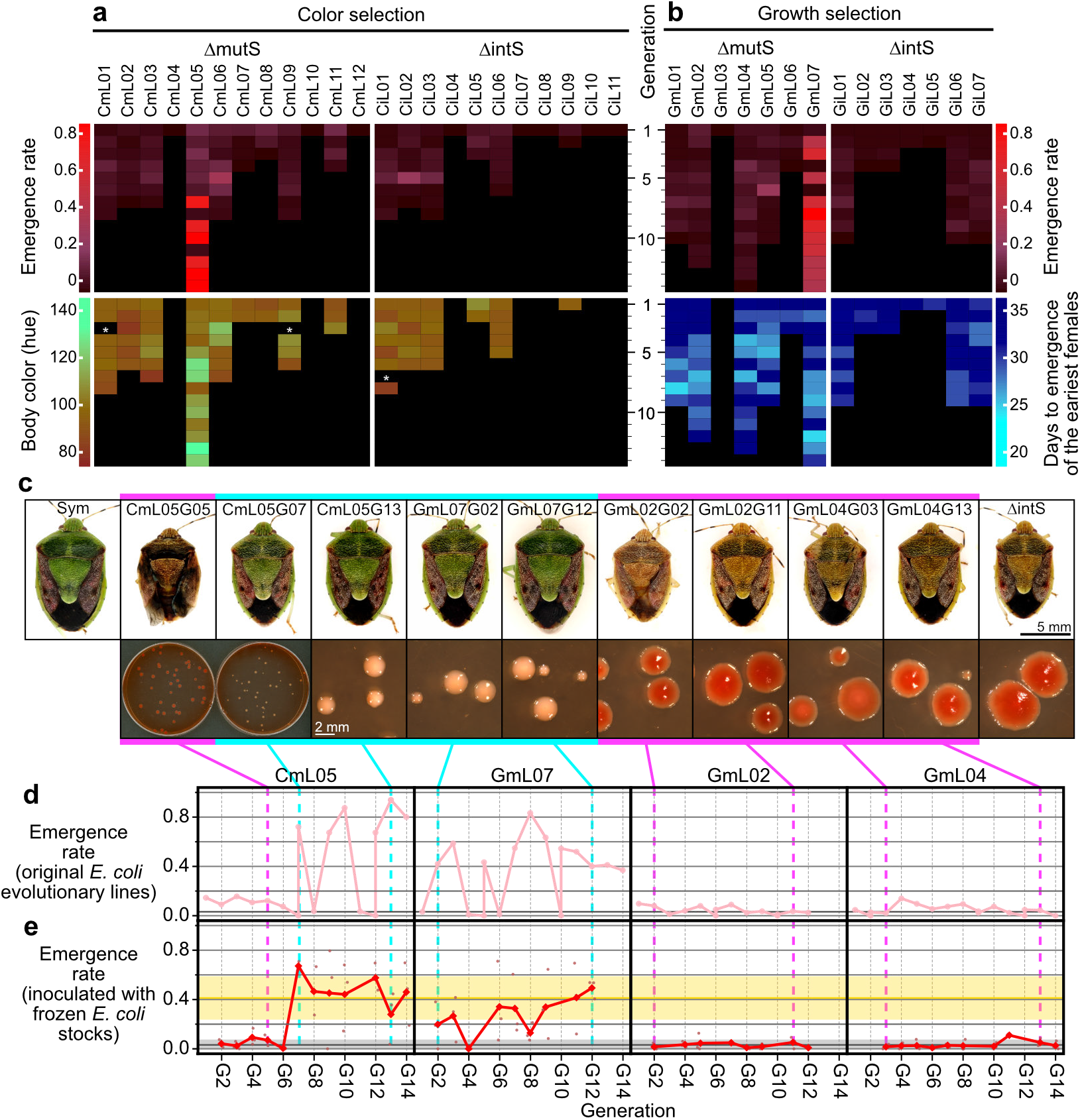
Evolution of mutualistic traits for *P. stali* in hyper-mutating *E. coli* lines. (**a**) Evolutionary *E. coli* lines subjected to host’s body color selection. Data of adult emergence rate and body color are displayed by heat maps. White asterisks indicate missing data of body color measurement. (**b**) Evolutionary *E. coli* lines subjected to host’s growth speed selection. Data of adult emergence rate and days to the first adult emergence are displayed by heat maps. Note that in (**a**) and (**b**), when an evolutionary line produced no adult insect and recovery from the freeze stock failed twice consecutively, the evolutionary line was terminated due to shortage of inoculum. From generation 10 and on, selected evolutionary lines were maintained. (**c**) Host’s body color and colony morphology of evolutionary *E. coli* lines. Red colonies are due to rich extracellular matrix produced on the agar plates containing Congo red. (**d, e**) Adult emergence patterns of *P. stali* infected with the representative *E. coli* lines, CmL05, GmL07, GmL02 and GmL04, in the original evolutionary experiments (**d**) and those in the confirmation experiments using frozen *E. coli* stocks (**e**). In (**c**)-(**e**), magenta lines and blue lines highlight “improved” generations and “non-improved” generations, respectively.

## Microbial traits of mutualistic *E. coli*

In addition to the colony size, shape and extracellular matrix on agar plates (Fig. 2c), the mutualistic *E. coli* lines CmL05 and GmL07 in culture exhibited distinct microbial traits in comparison with the original *E. coli* strains: slower growth rate, smaller cell size, loss of flagellar motility, and unstable cell shape (fig. S6a-g). Within the host insect, the evolutionary *E. coli* lines CmL05 and GmL07 showed significantly higher infection densities than the original *E. coli* strains (fig. S6h). These observations revealed that the mutualistic *E. coli* lines certainly have developed a variety of “symbiont-like” microbial traits.

## Transcriptomics and genomics of mutualistic *E. coli*

An aliquot of the dissected symbiotic organ from each generation of the color selection line CmL05 was subjected to RNA sequencing, from which *E. coli*-derived reads were extracted and analyzed (tables S1 and S2). Interestingly, the gene expression patterns of *E. coli* at generations 7-14 after the improvement of host phenotypes were separately clustered in contrast to those at generations 1-6 before the improvement (Fig. 3a). In the growth selection line GmL07, similarly, the gene expression patterns of *E. coli* at generations 2-12 after the improvement were distinct from that at generation 1 before the improvement and also from those of the other growth selection lines GmL02 and GmL04 in which the improvement of host phenotypes did not occur (Fig. 3b; tables S1 and S3). These results suggested that the evolution of the mutualistic *E. coli* lines entails specific and global change of gene expression patterns.

**Fig. 3.**
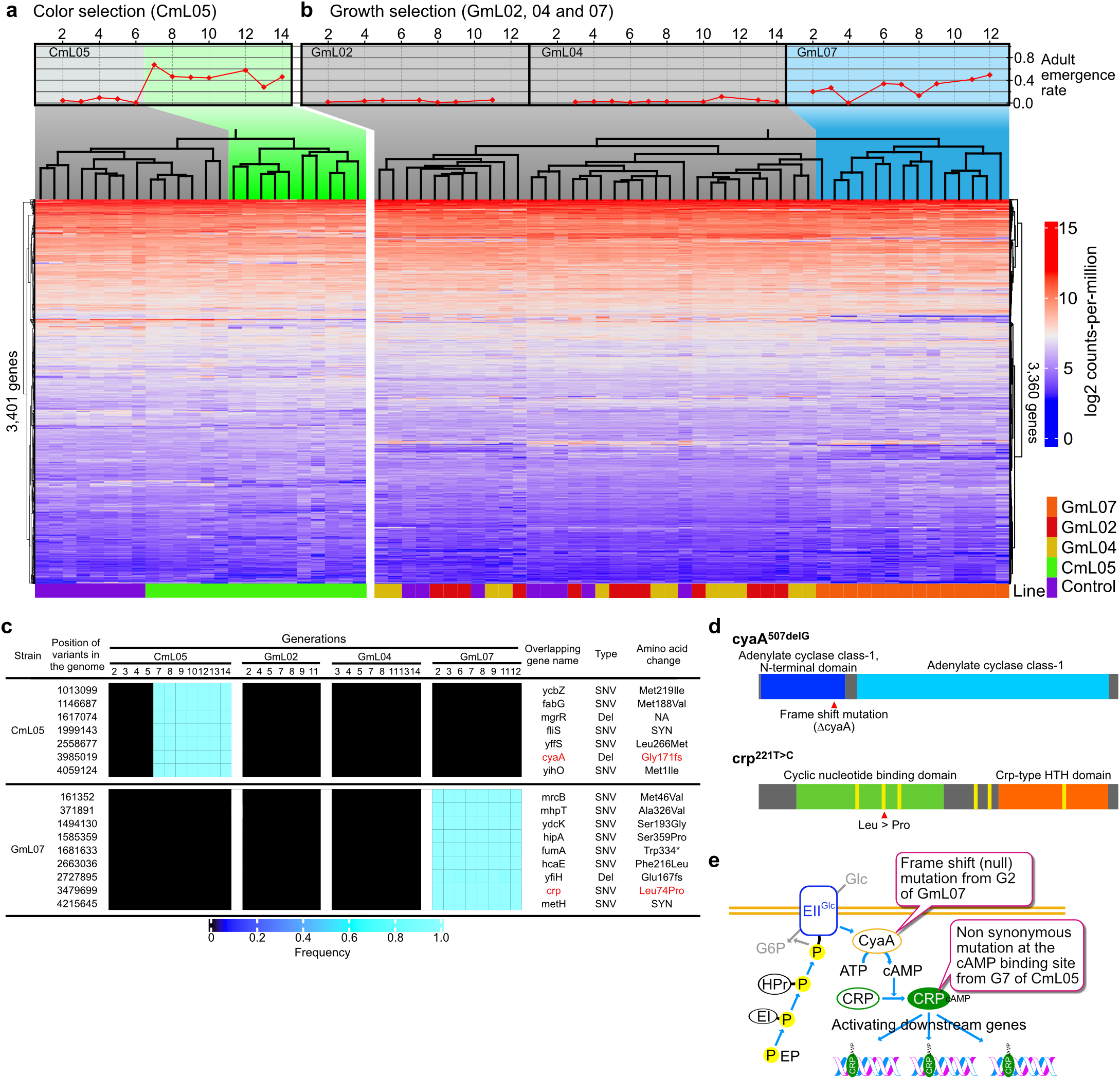
Transcriptomics and genomics of evolutionary *E. coli* lines. (**a, b**) Clustering dendrograms and heatmaps based on gene expression levels across generations of evolutionary *E. coli* lines subjected to color selection (right, 3,401 genes) (**a**) and growth selection (left, 3,360 genes) (**b**). Gray and colored areas depict non-improved and improved generations, respectively. (**c**) Mutations identified in the genomes of CmL05 and GmL07 as coincident with the improvement of host phenotypes. (**d**) Candidate mutations disrupting the catabolite repression pathway: a frame shift mutation in cyaA of CmL05 (top) and a non-synonymous mutation causing change from leucine to proline at a functionally important cAMP binding domain in crp of GmL07 (bottom). (**e**) Schematic presentation as to how CRP pathway is disrupted by the cyaA and crp mutations.

In the growth selection line GmL07 and the color selection line CmL05, we surveyed differentially expressed genes before and after the improvement of host phenotypes (table S4), which identified 193 commonly down-regulated genes and 95 commonly up-regulated genes across GmL07 and CmL05 (fig. S7a, b). The commonly down-regulated genes contained a number of metabolism-related genes such as transporter genes for sugars and other nutrients like maltose, ribose, galactitol, trehalose, mannose, branched chain amino acids, etc., glyoxylate bypass genes, fatty acid degradation genes, and others. Notably, core genes involved in extracellular matrix (= Curli fimbriae) production were significantly down-regulated after the improvement (fig. S7c), which accounted for the altered colony morphology of *E. coli* associated with the improvement of host phenotypes (see Fig. 2c).

The improved lines CmL05 and GmL07 and the non-improved lines GmL02 and GmL04 were subjected to genome sequencing throughout the evolutionary course (table S5), which identified many mutations accumulated in the hyper-mutating *E. coli* lines (tables S6 and S7; fig. S8). In an attempt to identify candidate mutations that are correlated with the improvement of the host phenotypes, we surveyed the mutations that appeared at generation 7 of CmL05 and then fixed, which yielded 7 candidate genes, and also the mutations that appeared at generation 2 of GmL07 and then fixed, which yielded 9 candidate genes (Fig. 3c).

## Disrupted CCR pathway in mutualistic *E. coli*

Of these candidates, we focused on a frame shift mutation that disrupted adenylate cyclase (CyaA) in CmL05, and a non-synonymous mutation that changed a functionally important cAMP binding site of cAMP receptor protein (Crp) from leucine to proline in GmL07 (Fig. 3d). Despite their independent origins in distinct evolutionary lines, CyaA and Crp are pivotal components of the same global metabolic regulator system, the carbon catabolite repression (CCR) pathway, operating in diverse bacteria including *E. coli* (*16,17*) (Fig. 3e). With sufficient availability of glucose as the primary carbon source for *E. coli,* the CCR components are subjected to glucose-mediated suppression, being in an unphosphorylated form incapable of activating CyaA, by which the intracellular cAMP is maintained at a low level (fig. S9a). When glucose is used up, the glucose-mediated suppression is released, by which the CCR components are phosphorylated and activate CyaA, which results in an elevated intracellular cAMP level and promotes cAMP binding to Crp. The resultant global transcriptional regulator Crp-cAMP activates and/or represses several hundreds of operons throughout the bacterial genome, referred to as the Crp-cAMP regulon, by which the bacterial metabolic pathways are switched to exploit other carbon sources for adaptation to nutrient-deficient and/or high bacterial density conditions (fig. S9b) (*18,19*). According to RegulonDB (*20*), the Crp-cAMP regulon of *E. coli* consists of some 390 up-regulated genes and 80 down-regulated genes (fig. S9c), which are involved in, for example, up-regulation of transporters and catabolic enzymes for non-glucose sugars (*21*), quorum sensing induction (*22*), and production of extracellular matrix (*23*).

Both the CyaA mutation in CmL05 and the Crp mutation in GmL07 are disruptive of the CCR pathway. Considering that *E. coli* cells are packed in the host symbiotic organ very densely (see Fig. 1k; fig. S2i, k), it seems likely that the symbiotic *E. coli* may be under a nutrient-limited condition within the host insect, at least locally. If so, it is expected that, in the evolutionary *E. coli* lines, while the Crp-cAMP transcriptional regulator was activated before the mutations occurred, the activation was disabled after the mutations occurred. Notably, of 193 genes commonly down-regulated after the CyaA mutation in CmL05 and the Crp mutation in GmL07, 55 genes were reported as activated by Crp-cAMP (fig. S10a). These genes, which are expected to be silenced upon disruption of the CCR system, were significantly down-regulated in CmL05 and GmL07, which represented many transporter genes for non-glucose sugars, carbohydrate metabolism genes, quorum sensing genes, extracellular matrix production genes, transcription factor genes, and others (fig. S10b-i).

## Disrupted CCR genes make *E. coli* an insect mutualist

In order to test whether these mutations are involved in the mutualistic traits of the evolutionary *E. coli* lines, we prepared *E.coli* strains that carry the mutations under the wild-type genetic background: the strain ΔcyaA in which cyaA gene is disrupted; and the strain crp^221T>C^ whose crp gene was engineered to carry the leucine-proline replacement at the cAMP binding site. Both the mutant *E. coli* strains exhibited small and convex colonies with little extracellular matrix, somewhat slower growth rate, smaller cell size, and loss of flagellar motility (Fig. 4a; fig. S11a-e), which were generally reminiscent of the characteristic traits of the improved evolutionary *E. coli* lines CmL05 and GmL07 (Fig. 2c; fig. S6a-e). When the mutant *E. coli* strains were inoculated to sterilized newborn nymphs of *P. stali*, both the ΔcyaA-infected insects and the crp^221T>C^-infected insects exhibited remarkably high adult emergence rates, which were comparable to the insects infected with the improved evolutionary *E. coli* lines and were significantly higher than the insects infected with the control *E. coli* strains (Fig. 4b). Moreover, the ΔcyaA-infected insects and the crp^221T>C^-infected insects were greenish in color, which were comparable to the greenish insects infected with the improved evolutionary *E. coli* lines and distinct from the dwarf brown insects infected with the control *E. coli* strains (Fig. 4c). On the other hand, infection densities of crp^221T>C^ and ΔcyaA were not comparable to those of the improved evolutionary *E. coli* lines (fig. S11f). These results demonstrated that, strikingly, the single mutations that disrupt the CCR global regulator system make *E. coli* mutualistic to the host insect *P. stali*.

**Fig. 4.**
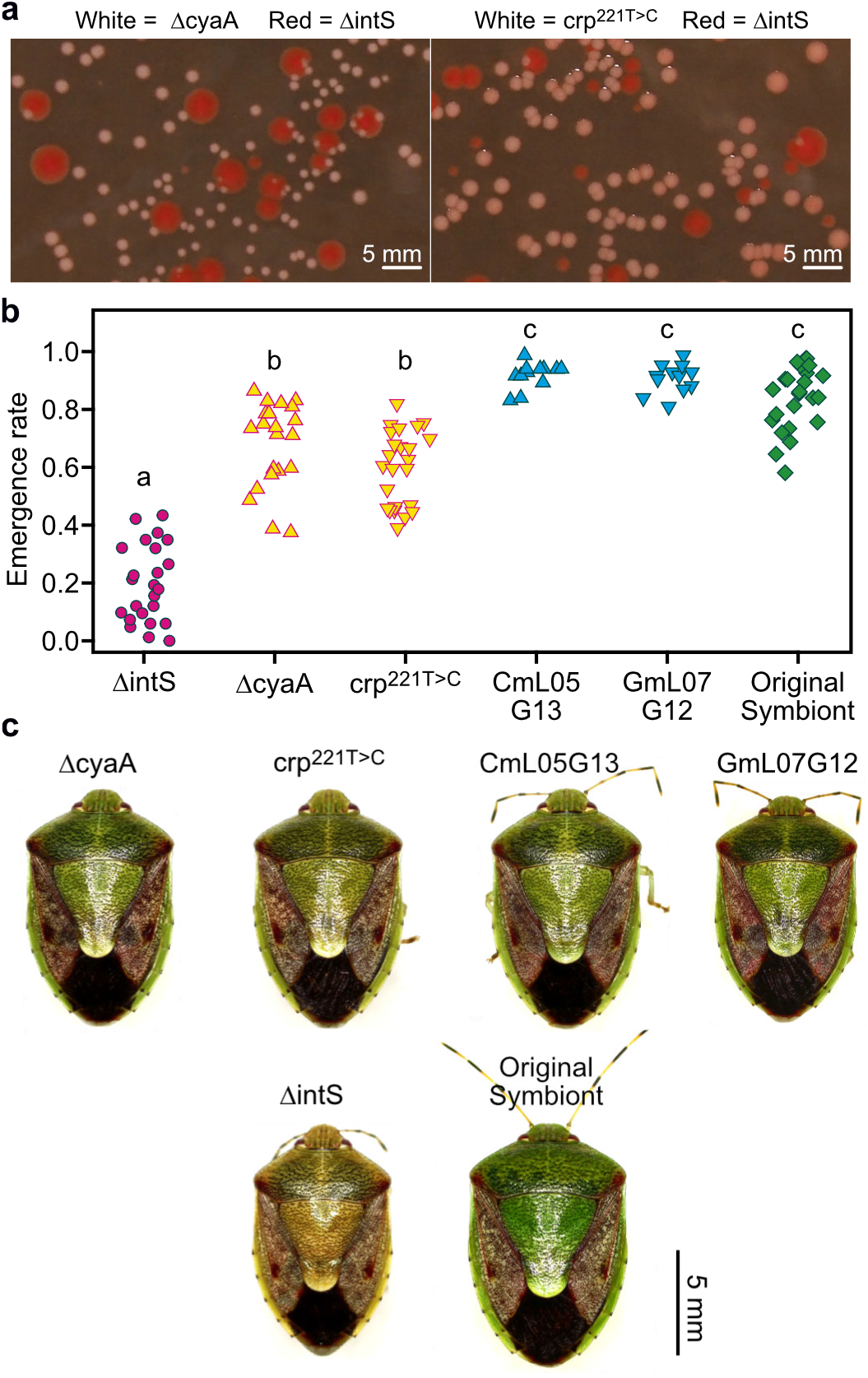
Single mutations disrupting carbon catabolite repression pathway make *E. coli* mutualistic to *P. stali.* (**a**) Small, convex and white colonies of ΔcyaA and crp^221T>C^. (**b**) Adult emergence rates of *P. stali* infected with ΔcyaA and crp^221T^>^C^. Different alphabetical letters indicate statistically significant differences (pairwise Wilcoxon rank sum test with Bonferroni correction: *P* < 0.05). (**c**) Adult insects infected with ΔcyaA and crp^221T>C^, which are larger in size and green in color in comparison with those infected with control ΔintS.

## Discussion

We established an experimental insect-*E. coli* symbiotic system in which the model bacterium is localized to host symbiotic organ, transmissible to host offspring vertically, and supportive of host survival, though incompletely. By infecting and passaging a hyper-mutating *E. coli* strain with the host insect trans-generationally, several evolutionary lines rapidly developed improved adult emergence and body color, realizing recurrent evolution of mutualism in the laboratory. Strikingly, the *E. colis* evolution into the insect mutualist was ascribed to single mutations that convergently disrupted the bacterial CCR pathway, uncovering unexpected involvement of the nutrient-responsive global transcriptional regulator in the establishment of symbiosis.

Our finding sheds new light on the evolvability of symbiosis – elaborate mutualistic symbiosis can evolve much more easily and rapidly than conventionally envisaged. We suggest the possibility that the inactivation of the CCR global regulator may represent a pivotal evolutionary step at an early stage of symbiosis. Densely packed in the symbiotic organ, the symbiotic bacteria are expected to constantly suffer nutritional shortage and activate the CCR pathway in vain, which may incur substantial metabolic cost and destabilize the symbiotic association. In this context, the disruption of the CCR pathway should benefit and stabilize the symbiosis. Our finding may be also relevant to the general evolutionary trend of symbiont genomes toward size reduction (*24*) and lack of transcription factors (*25*). The disruption of the CCR pathway causes silencing of otherwise activated about 400 genes under the Crp-cAMP regulon (*20*), which accounts for about 10% of the whole *E. coli* genome and provides potential targets for gene disruption, IS bombardment, intragenomic recombination, and reductive genome evolution. We propose that, although speculative, inactivation of transcriptional regulators and genome size reduction might have concurrently proceeded in this way during the symbiont genome evolution.

The *P. stali-E. coli* experimental symbiotic system will open a new window to directly observe and analyze the evolutionary processes and mechanisms of mutualistic symbiosis in real-time. *E. coli* is among the best understood cellular organisms, whose 4.5-5.5 Mb genome encodes over 4,000 genes and around 70% of them are with functional information (*26,27*). Laboratory evolution of mutualism using such a model bacterium with ample technological and genetic resources will lead to an ultimate understanding of the symbiotic evolution. Considering that *E. coli* represents a universal component of the gut microbiome of human, mouse, and other vertebrates (*28*), the insect-*E. coli* system in combination with the germfree mouse-*E. coli* experimental evolution systems (*29,30*) would enable us to pursue not only the differences but also the commonality underpinning the mechanisms of gut symbiosis across vertebrates and invertebrates.

## Methods

### Insect and bacterial strains used in this study

An inbred laboratory strain of the brown-winged green stinkbug *P. stali* was established from several adult insects collected at Tsukuba, Ibaraki, Japan in September 2012, and has been maintained in the laboratory for years. This strain is associated with an essential and uncultivable gut symbiont *Pantoea* sp. A (*11*) in a posterior midgut region specialized as the symbiotic organ (Fig. 1; fig. S2). The insects were reared on raw peanuts, soybeans and water containing 0.05% ascorbic acid (Merck, Germany) at 25 ± 1 °C and 50 ± 5% relative humidity under a long-day regime of 16 h light and 8 h dark.

*E. coli* strains and mutants used in this study are listed below. The mutants ΔintS, ΔmutS and crp22iT>c were generated as described later.

| Name | Description | Reference |
| --- | --- | --- |
| BW25113 | Obtained from National Bio Resource Project (NBRP) | 31 |
| ΔintS | The intS gene was replaced with kanamycin resistance gene. | This study |
| ΔmutS | The mutS gene was removed from ΔintS. | This study |
| ΔcyaA | Obtained from Keio single-gene knock-out mutant library via NBRP | 27 |
| $Crp^{221T>C}$ | A nonsynonymous single nucleotide substitution was introduced in the *crp* gene | This study |
| CmL05G13a | An *E. coli* clone isolated from the glycerol stock of CmL05G13. | This study |
| GmL07G12a | An *E. coli* clone isolated from the glycerol stock of GmL07G12. | This study |
| EPI300 | Obtained from Epicentre | N.A. |
| DH5α | Common laboratory strain | N.A. |
| JM109 | Common laboratory strain | N.A. |
| BL21 | Common laboratory strain | N.A. |

### Construction of *E. coli* mutants

The *E. coli* mutant ΔintS was established by replacing intS gene of *E. coli* BW25113 with nptII gene that confers kanamycin resistance by λ-Red homologous recombination using the pRed/ET plasmid (Gene Bridges, Germany). The *E. coli* mutant ΔmutS was established by replacing mutS gene of ΔintS with the FRT-Cm-FRT cassette (Gene Bridges, Germany) by λ-Red homologous recombination, and then Cm^R^ was eliminated by Flp-FRT recombination. The *E. coli* mutant crp^221T>C^ was established from ΔintS by replacing the 221^st^ nucleotide T of the wild type crp gene with C, which changed 74^th^ amino acid leucine of the Crp protein to proline. This replacement was introduced by MAGE method (*32*) with a 90-mer DNA oligo (5’-taaagaaatg-atcctctcct-atctgaatca-gggtgatttt-attggcgaac-<u>**C**</u>gggcctgtt-tgaagagggc-caggaacgta-gcgcatgggt) whose 1^st^ to 4^th^ nucleotides were phosphothioated.

### Preparation of symbiont-free nymphs by surface sterilization of eggs

Egg clutches produced by the stock culture of *P. stali* were soaked in 4% formaldehyde for 10 min, rinsed with sterilized water several times, and kept in sterilized plastic boxes until use. While this treatment does not affect hatchability and survival of the eggs, newborn nymphs fail to acquire the symbiotic bacteria and become symbiont-free (*33*).

### Experimental evolution of *P. stali–E. coli* artificial symbiotic system

Evolutionary experiments in this study consisted of, for each evolutionary *P. stali* line, (i) preparation of an inoculum either from *E. coli* culture of ΔmutS or ΔintS (only G1) or from an adult female of the previous generation (from G2 and on), (ii) oral administration of the inoculum to symbiont-free nymphs, (iii) rearing of the nymphs either to their adulthood or death, (iv) selection of an adult female for inoculation to the next generation, (iv) contamination check of the selected adult female, (v) preparation of an inoculum and a glycerol stock from the symbiotic organ dissected from the selected female, and (vi) morphological measurements of all adult insects obtained.

Either diluted *E. coli* culture (2.5 ml adjusted to OD_600_ = 0.1) or homogenate of the symbiotic organ dissected from a selected female of the previous generation (2.5 ml containing 1/2 organ equivalent) was soaked in a cotton pad and orally administered to around 84 symbiont-free hatchlings derived from six surface-sterilized egg masses, by making use of the nymphal behavior that, after egg surface probing for about 30 min and resting for around a day, they take water without feeding and molt to second instar in a few days (*11,33*). These nymphs were reared on sterilized peanuts, soybeans and ascorbic acid water as described previously (*33*). In the evolutionary experiments, two selection schemes, growth selection and color selection, were conducted (fig. S3). In growth selection lines (GmL for hyper-mutating ΔmutS lines; GiL for non-mutating ΔintS lines), the first-emerged adult female was subjected to dissection of the symbiotic organ for inoculation to the next generation as well as freeze storing. In color selection lines (CmL for ΔmutS lines; CiL for ΔintS lines), adult females were collected for 35 days after hatching or until at least one adult female emerged. These adult females were anesthetized on ice and photographed from the ventral side using a digital camera. Their body color was measured using the image analyzing software Natsumushi ver. 1.10 (*34*), and the adult female that exhibited the highest hue angle (= greenness) was subjected to dissection of the symbiotic organ for inoculation to the next generation as well as freeze storing.

The symbiotic organ of the selected female was dissected in PBS (0.8% NaCl, 0.02% KCl, 0.115% Na2HPO4, 0.02% KH2PO4, pH 7.4), rinsed with 70% ethanol, and homogenized in 200 μL sterile water. Of the 200 μL homogenate, 5 μL was used for contamination check by quantitative PCR. The number of *E. coli* genome copies was evaluated in terms of kanamycin resistance gene copies, which is present in the ΔintS and ΔmutS mutants but absent in wildtype *E. coli* and other bacteria. The number of total bacterial genome copies was evaluated based on bacterial 16S rRNA gene copies. When the former *E. coli* genome copy number was approximately the same as the latter bacterial genome copy number, the specimen was diagnosed as free of contamination. When the specimen was diagnosed as contaminated, the next best female was used. For quantitative PCR, the primers Tn5-1789F (5’-TGC TCG ACG TTG TCA CTG AA-3’) and Tn5-1879R (5’-GCA GGA GCA AGG TGA GAT GA-3’) were used for kanamycin resistance gene, while the primers 16S-967F (5’-CAA CGC GAA GAA CCT TAC C-3’) and 16S-1046R (5’-CGA CAG CCA TGC ANC ACC T-3’) were used for bacterial 16S rRNA gene. The PCR reaction was performed using Brilliant PCR mix (Agilent Technologies, USA). The standard curve was drawn using serially diluted ΔintS genomic DNA, which contains one kanamycin gene copy and seven 16S rRNA gene copies per genome. The thermal profile was the initial denaturation at 95°C for 3 min followed by 40 cycles of incubation at 95°C for 5 sec and at 60°C for 10 sec. To confirm specific amplification, melting curve analysis was also included. The reaction was conducted on Mx3000p (Agilent Technologies, USA). While 100 μL of the homogenate of the female symbiotic organ diagnosed as free of contamination was used as the inoculum to the next generation, the remaining homogenate (~95 μL) was mixed with an equal volume of 20% glycerol and stored at −80°C.

### Inoculation of *E. coli* frozen stocks to *P. stali*

The frozen glycerol stocks were thawed, of which 50 mL was taken and diluted with sterile water to 3 mL. Each of three replicates of around 84 symbiont-free hatchlings from six surface-sterilized egg masses was fed with 1 ml inoculum soaked in a cotton pad as described above. The symbiont A and ΔmutS were included in the evaluation as positive and negative controls, respectively. Adult emergence of the insects was monitored for 50 days after hatching. All the adult insects were photographed from the dorsal side with a digital camera, and hue angle (= greenness) of the scutellum and thorax width were measured using ImageJ (*35*). For the subsequent RNA sequencing analyses and resequencing of *E. coli* genomes, the symbiotic organs were isolated from the adult insects and homogenized in 100 μL PBS. Of 100 μL homogenate, 50 μL was subjected to RNA sequencing and the remaining 50 μL was used for genome resequencing.

### RNA sequencing analyses

The homogenate of the symbiotic organ was subjected to total RNA extraction using RNAiso (Takara Bio, Japan) and RNeasy Mini Kit (Qiagen, Nederland). Then, ribosomal RNAs of both insect and bacterial origins were removed from the total RNA samples using Ribo-Zero Gold rRNA Removal Kit (Epidemiology) (Illumina, USA). The rRNA-depleted RNAs were converted to paired end libraries using Sure Select Strand Specific RNA Kit (Agilent Technologies, USA) or TruSeq RNA Library Prep Kit v2 (Illumina, USA) (see table S1). The libraries were sequenced with Hiseq 3000 or Hiseq X (Illumina, USA).

The obtained sequences were trimmed, mapped to *E. coli* BW25113 genome sequence (Accession number NZ_CP009273), and read-counted with CLC Genomics Workbench 10.0 (Qiagen, Germany). Normalizations and differential expression analyses were conducted with EdgeR ver. 3.32.1 (*36*). Complex Heatmap ver. 2.10.0 (*37*) was used for clustering analyses and drawing heatmaps of the RNA sequencing libraries.

### Genome resequencing and detection of structural changes

DNA samples were extracted from the homogenates of the symbiotic organ using QIAamp DNA Mini Kit (Qiagen, Germany). The extracted DNAs were converted to paired end libraries using Nextera XT DNA Library Prep Kit (Illumina, USA) and the libraries were sequenced with Miseq (Illumina, USA). CLC Genomic Workbench ver. 10.0 was used for detection of *E. coli* genome variants that emerged during the evolutionary experiments. The heatmaps of the variant frequency data were drawn using Complex Heatmap (*37*).

### Fluorescence in situ hybridization

Fluorescence in situ hybridization (FISH) analyses were performed essentially as described (*38*). The whole insect bodies or isolated digestive tracts were fixed with PBS containing 4% formaldehyde (Fujifilm, Japan). The fixed samples were embedded in Technovit 8100 (Kulzer, Germany) and processed into 2 μm tissue sections using a rotary microtome RM2255 (Leica, Germany). The AlexaFluor555-labeled oligonucleotide probes Eco934 (5’-CAT GCT CCA CCG CTT GTG-3’) and SymAC89R (5’-GCA AGC TCT TCT GTG CTG CC-3’) were used to detect *E. coli* and the symbiont A, respectively (*12*). Host nuclei were counterstained with 4’, 6-diamidino-2-phenylindole (DAPI) (Dojindo, Japan). The hybridized specimens were observed using a fluorescence dissection microscope M165FC (Leica, Germany), an epifluorescence microscope DM6B (Leica, Germany), and a laser confocal microscope LSM700 (Zeiss, Germany).

### Infection of *E. coli* mutants and effects on host phenotypes

*E. coli* mutants were cultured, diluted, and orally administrated to symbiont-free newborn nymphs of *P. stali* as described above. The insects were reared to monitor their adult emergence for 42 days after hatching. The dorsal images of the adults were taken with an image scanner GT-X830 (Epson, Japan), and the hue angle of the scutellum and thorax width were measured and analyzed using the software Natsumushi (*34*). *P. stali* harboring the original symbiont *Pantoea* sp. A was also included as a reference. As for the adult females infected with *E. coli,* bacterial titers in the symbiotic organs were measured by quantitative PCR. KAPA SYBR Fast qPCR Kit (Roche, USA), Tn5-1789F and Tn5-1879R primer sets were used for quantification. The standard curves were drawn using serially diluted pT7Blue (Takara Bio, Japan) plasmid carrying a kanamycin resistance gene fragment. The quantitative PCR reactions were conducted on Light Cycler 96 (Roche, Switzerland).

### Measurement of *E. coli* phenotypes

For inspection of colony morphology and extracellular matrix production, *E. coli* cultures were spread onto LB agar plates containing 80 μg/mL Congo Red (Merck, USA) and incubated at 25°C for 3 days. Colonies formed on the plate were photographed by using a scanner GT-X850 and/or dissection microscope S9i (Leica, Germany).

For growth curve measurements, each glycerol stock of *E. coli* was inoculated to 2 mL LB broth (Becton Dickinson, USA) and incubated at 25°C for 16 h with shaking at 200 rpm. The cell culture was diluted to OD_600_ = 0.005 in 25 mL LB broth, and incubated at 25°C with shaking at 200 rpm. From the bacterial culture, 120 μL of cell suspension was sampled every hour, and the samples were subjected to measurement of OD_600_ using a spectrometer UV-1800 (Shimadzu, Japan).

For time-lapse analyses of growth and morphology of individual *E. coli* cells, two types of microfluidic devices were used. One type was a microfluidic device in which bacterial cells were enclosed in microchambers etched on a glass coverslip. A cellulose membrane was attached to a coverslip via biotin-streptavidin binding, on which the microchambers were fabricated as described previously (*39,40*). Another type was a microfluidic device made of polydimethylolefin (PDMS) with a channel structure similar to Mother Machine 3 as described previously (*41*) (see fig. S6f). The width of the cell observation channels in this device was 9 μm, which was broader than that of the Mother Machine and thus each cell observation channel could harbor 30-70 individual *E. coli* cells depending on cell sizes. *E. coli* cells in exponential phase were introduced into both types of the microfluidic devices and observed under a Nikon Ti-E microscope (Nikon, Japan) equipped with ORCA-fusion camera (Hamamatsu Photonics, Japan). In the time-lapse measurements, phase-contrast images were acquired with a 100 × oil immersion objective lens (Plan Apol, NA 1.45) at an interval of 3 min, in which 50-100 XY positions were simultaneously observed. The microscope was controlled from a computer using Micromanager 4. In the microchamber device measurements, LB broth was supplemented with 0.1% bovine serum albumin and 0.02% Tween-80 to suppress cell adhesion, and introduced into the devices at a flow rate of 2 mL/h.

For measurements of size and flagellar motility, *E. coli* cells were grown in LB medium with shaking at 25°C to around OD_600_ = 2.0, observed under a phase-contrast microscope IX71 (Olympus, Japan), recorded by a CCD camera DMK33UP5000.WG (The Imaging Source, Germany) at 30 frames per second, and analyzed using ImageJ v1.53 (*35*) and IGOR Pro 8.02 J (WaveMetrics, USA). The cell size data were measured for individual six cultures. The swimming ratio data were obtained as the number of swimming cells in 100 cells from individual eight cultures.

### Statistics

Statistical analyses were conducted by using R ver. 4.1.2 (*42*) and RStudio (*43*). R was also used to plot the data.

## Acknowledgments

We thank U. Asaga, S. Kimura, J. Makino and T. Matsushita for insect rearing and technical assistance. This study was supported by the JST ERATO grants JPMJER1803 and JPMJER1902 (TF, CF, YW, RK) and the JSPS KAKENHI grant JP25221107 (TF, RK). Genome sequencing and analyses were supported by the JSPS KAKENHI grant JP16H06279.

## Author contributions

RK and TF conceived the project and designed the experiments. RK, MMo, NOT and YN performed insect-*E. coli* evolutionary experiments, RK, MMo, NOT, YI, HT, YN and THo analyzed insect phenotypes, RK, MMi, KO, RO and YW analyzed *E. coli* phenotypes, RK, MMo, NOT, YG and THa performed genome sequencing and analyses, MMo, RK, NOT, MS and YS conducted RNA sequencing and analyses, RK, HT, SS and CF designed and generated hyper-mutating and other *E. coli* strains, and TF wrote the manuscript with input from all the other authors.

## Competing interests

The authors declare no competing interests.

## Data and materials availability

All RNA sequencing and DNA sequencing data produced in this study were deposited in DDBJ Sequence Read Archive (DRA) (see tables S1 and S5). All data are available in the manuscript or the supplementary materials.

---

**Fig. S1.**
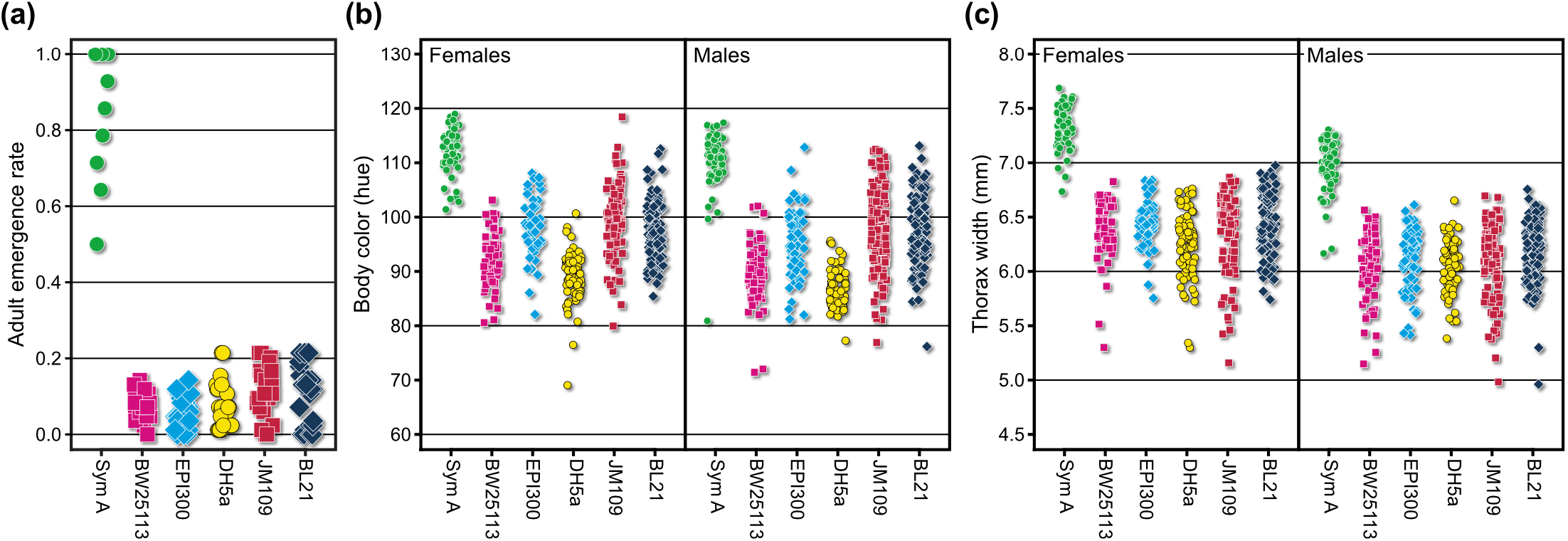
Phenotypes of *P. stali* adults infected with laboratory strains of *E. coli.* (**a**) Adult emergence rate. (**b**) Body color (greenish hue) of females (left) and males (right). (**c**) Body size (thorax width) of females (left) and males (right). Sym A is *Pantoea* sp. A, the original, uncultivable and essential gut symbiont of *P. stali* (*11*). BW25113, EPI300, DH5a, JM109 and BL21 are commonly used laboratory strains of *E. coli*.

**Fig. S2.**
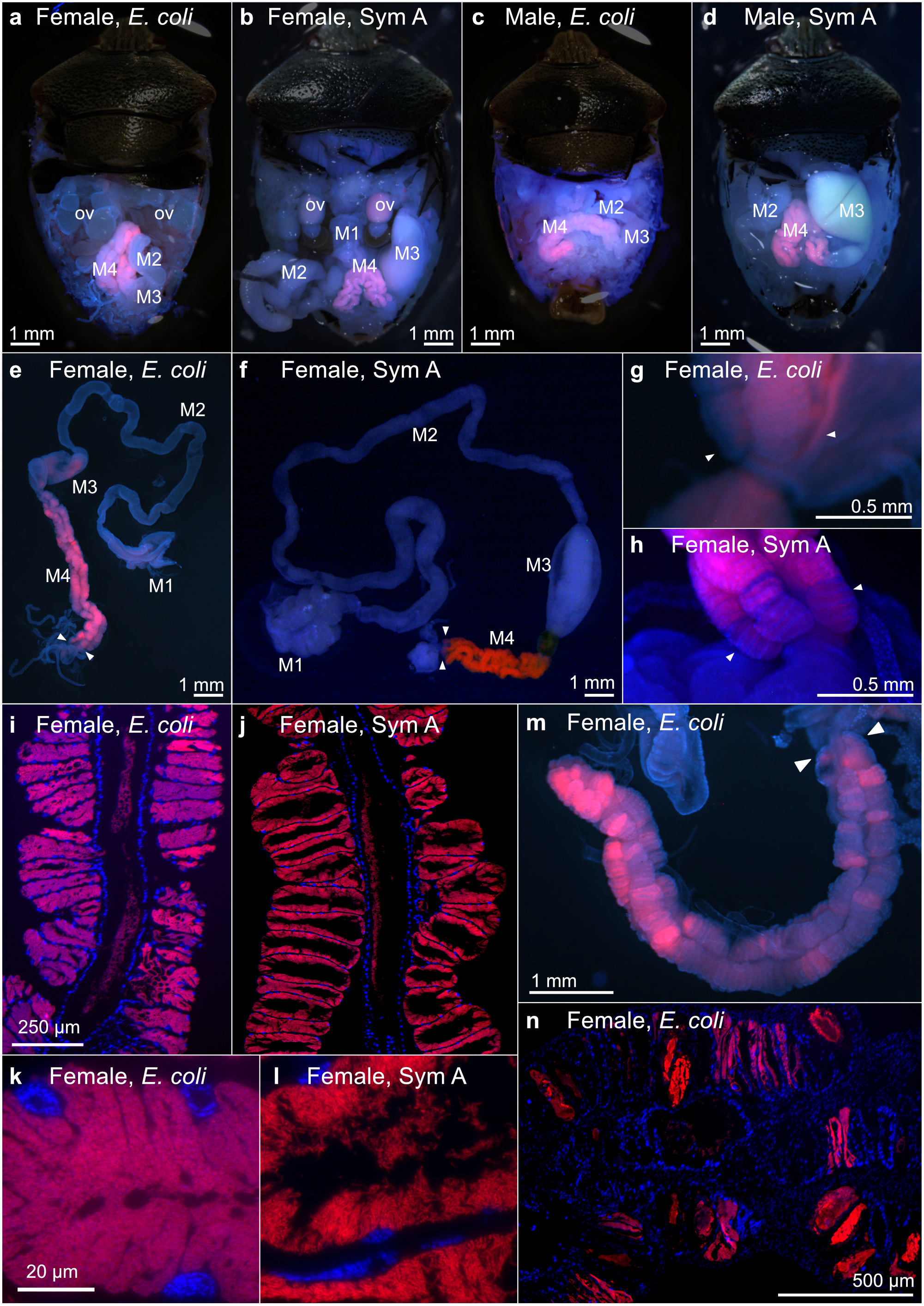
FISH localization of *E. coli* and original symbiont *Pantoea* sp. A (= Sym A) in *P. stali*. (**a-d**) Localization in abdominal body cavity of adult insects: (**a**) *E. coli* in adult female, (**b**) Sym A in adult female, (**c**) *E. coli* in adult male, and (**d**) Sym A in adult male. FISH signals are localized to the midgut M4 region. Signals in oocytes are due to autofluorescence. Abbreviations: M1, M2, M3, and M4, midgut regions M1, M2, M3, and M4 (= symbiotic organ); ov, ovary. (**e, f**) Localization of *E. coli* (**e**) and Sym A (**f**) in dissected alimentary tract of adult females. Arrowheads indicate female-specific enlarged end crypts at the posterior end of the symbiotic organ, which are presumably involved in vertical symbiont transmission by storing bacteria-containing secretion (*44*). (**g, h**) Magnified images of the end crypts infected with *E. coli* (**g**) and Sym A (**h**). Note that *E. coli*-infected end crypts are atrophied in comparison with Sym A-infected ones. (**i, j**) Localization of *E. coli* (**i**) and Sym A (**j**) in the crypt cavities of the symbiotic organ. (**k, l**) Magnified images of *E. coli* cells (**k**) and Sym A cells (**l**) packed in the crypt cavity. (**m, n**) Patchy localization patterns of *E. coli* in the symbiotic organ, which are often found with *E. coli* but seldom observed with Sym A.

**Fig. S3.**
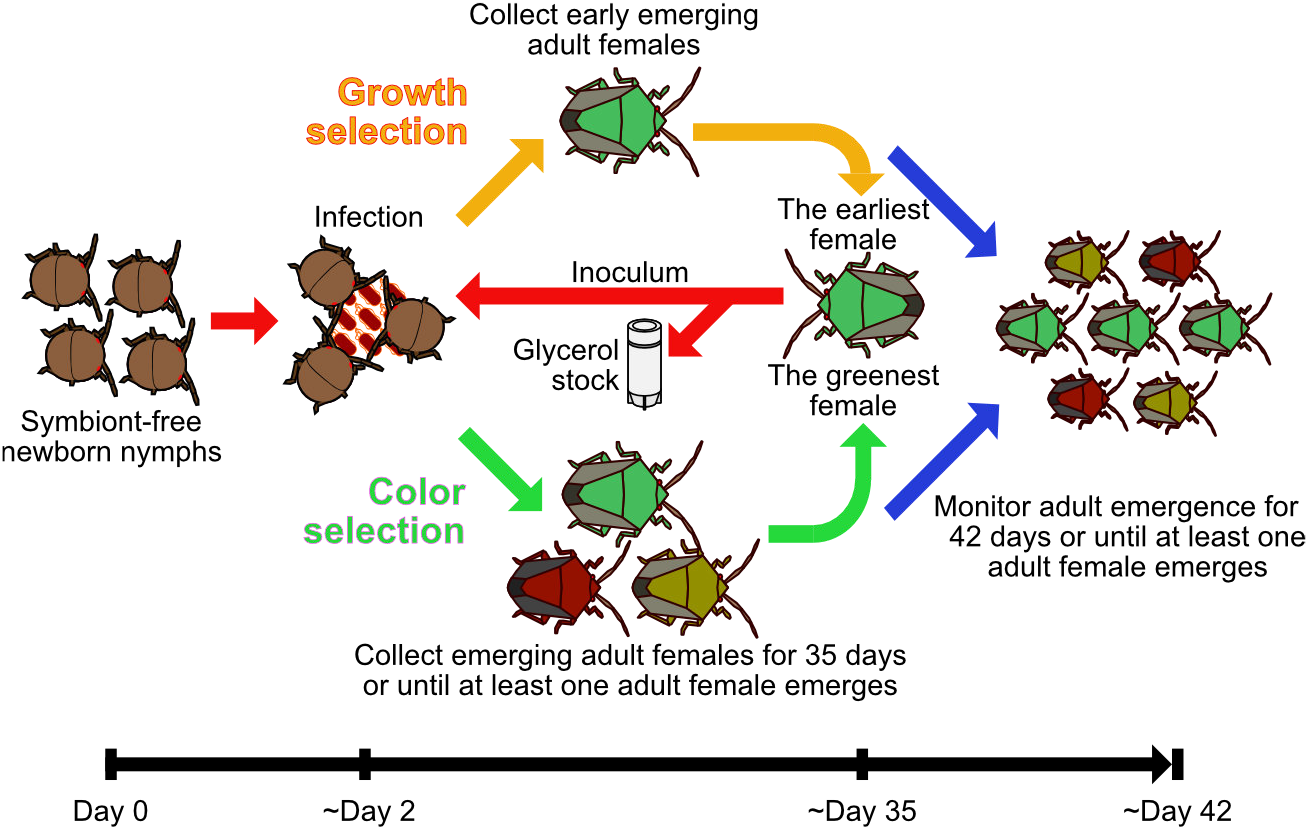
Experimental scheme for evolution of mutualistic *E. coli* with *P. stali.*

**Fig. S4.**
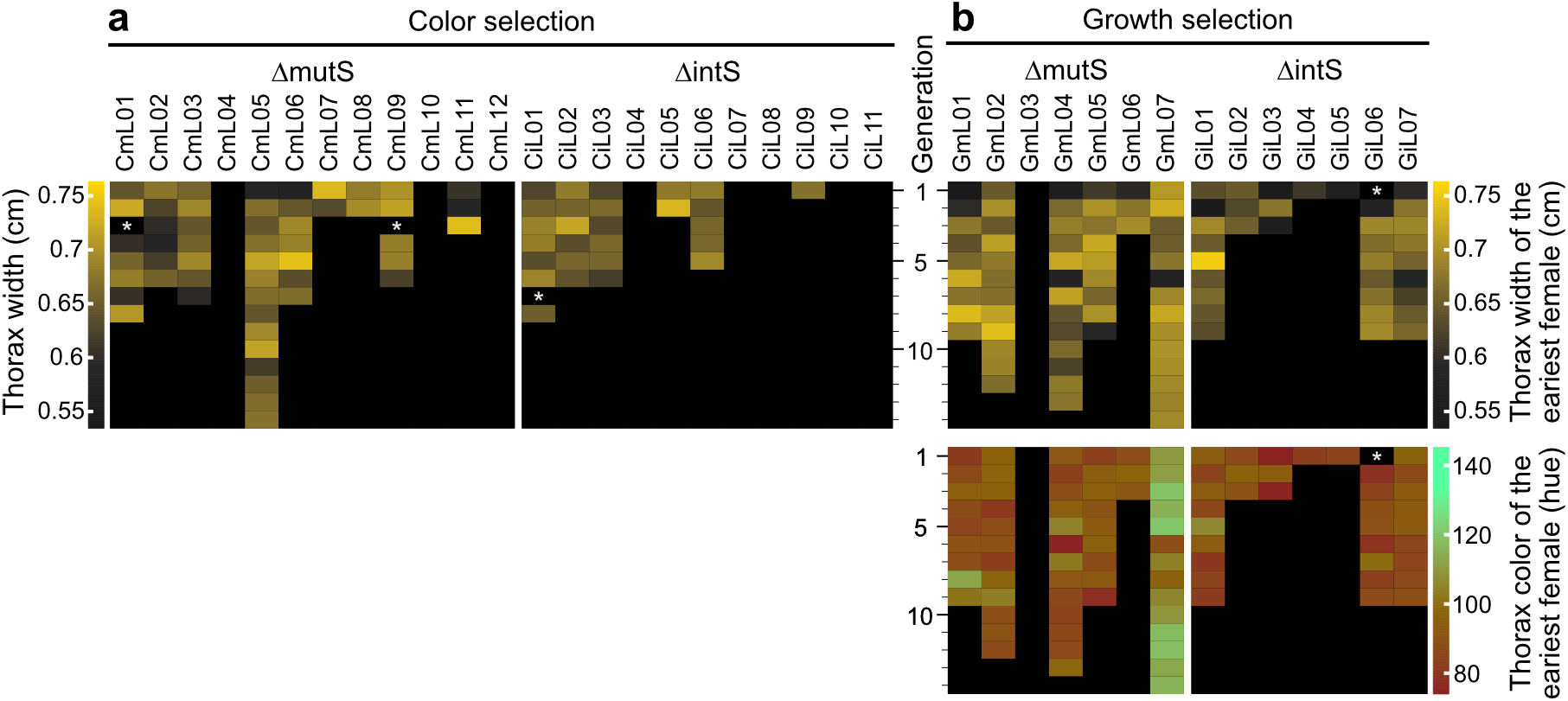
Effects of evolutionary *E. coli* lines on body size and color of *P. stali.* (**a**) Evolutionary *E. coli* lines subjected to host’s body color selection. Data of host’s body width are displayed by heat maps. Also see Fig. 2a. (**b**) Evolutionary *E. coli* lines subjected to host’s growth speed selection. Data of host’s body width and color are displayed by heat maps. Also see Fig. 2b.

**Fig. S5.**
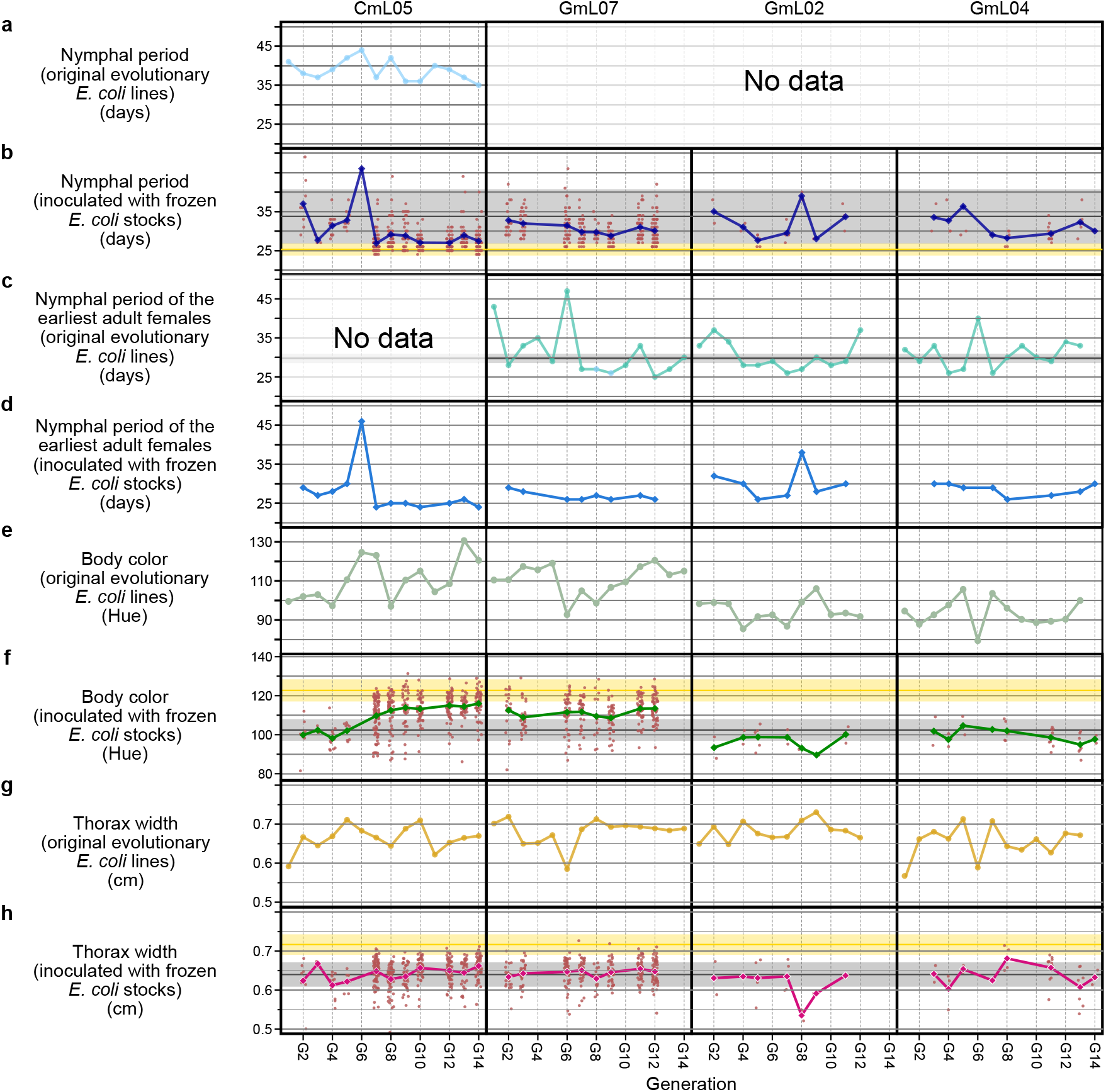
Adult phenotypes of *P. stali* infected with the evolutionary *E. coli* lines CmL05, GmL07, GmL02 and GmL04. (**a, b**) Nymphal period. (**c, d**) Nymphal period of the earliest adult females. (**e, f**) Body color. (**g, h**) Thorax width. (**a, c, e, g**) Phenotypes of adult insects infected with the original evolutionary *E. coli* lines. (**b, d, f, h**) Phenotypes of adult insects inoculated with the frozen *E. coli* stocks. Line charts show mean values while dots indicate individual data points. Note that, corresponding to each original evolutionary *E. coli* line, three insect groups were inoculated with the frozen *E. coli* stock.

**Fig. S6.**
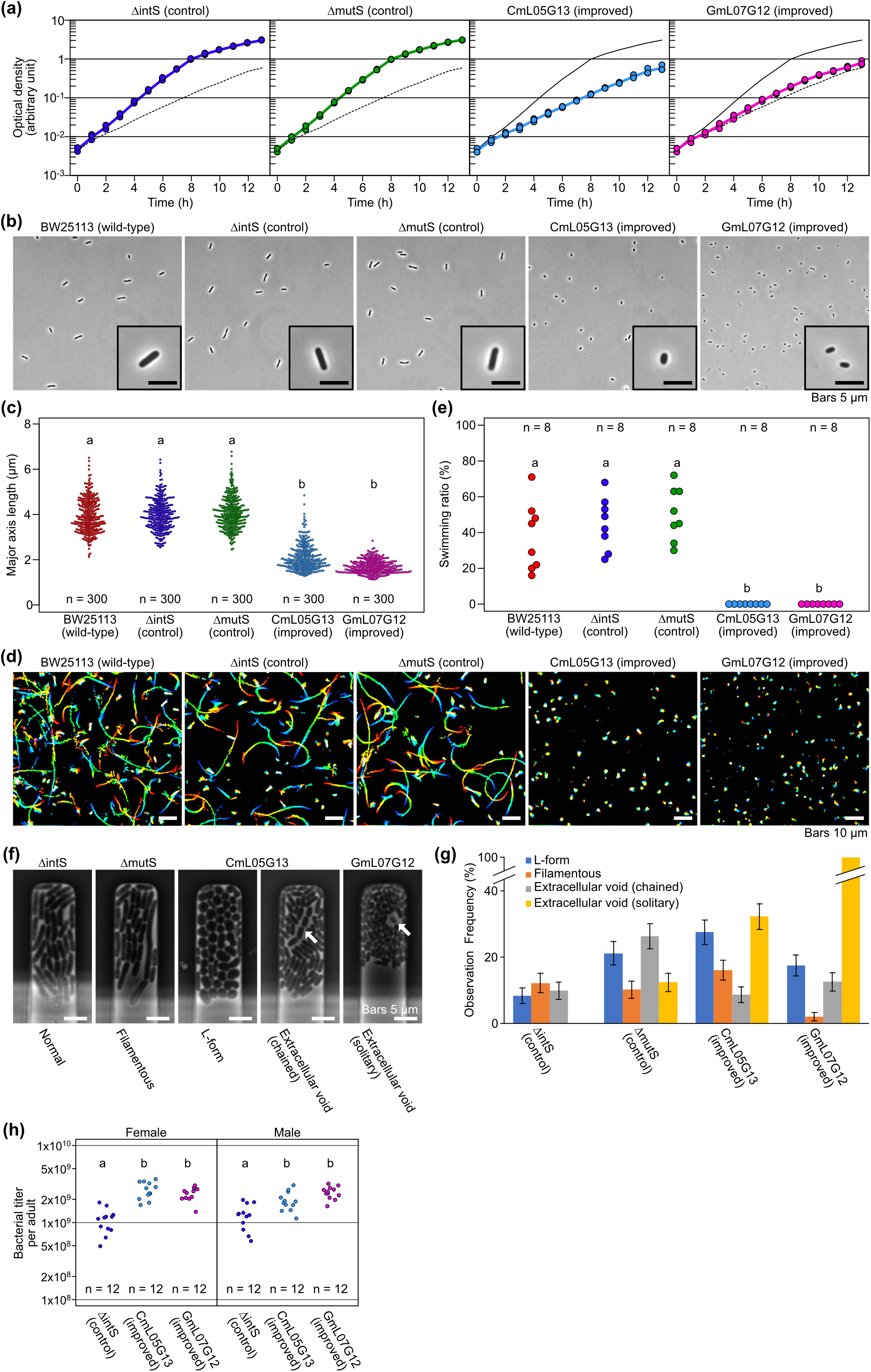
Microbial traits of evolutionary *E. coli* lines CmL05 and GmL07 in comparison with original *E. coli* strains BW25113, ΔintS and ΔmutS cultured in liquid medium. (**a**) Growth curves (3 replicates each). Upper solid line is the trace of ΔintS growth curve, whereas lower dotted line is the trace of CmL05 growth curve. (**b**) Morphology of bacterial cells. (**c**) Quantification of cell size in terms of major axis length. (**d**) Motility of bacterial cells visualized by rainbow plot for 2 sec. (**e**) Quantification of bacterial motility in terms of number of swimming cells per 100 cells observed. (**f**) Characteristic cellular shape and growth mode in microfluidic channels. From left to right, the micrographs show the microchannels harboring *E. coli* cells with normal rod-like shape (ΔintS), filamentation shape (ΔmutS), L-form-like round shape (CmL05), extracellular void space and chained growth (CmL05), and extracellular void space and solitary growth (GmL07). Arrows indicate the cells showing the extracellular void space. (**g**) Frequency of the microchannels in which *E. coli* cells exhibited characteristic cell shape and growth mode. The total numbers of microchannels observed in the time-lapse measurements were 131 (ΔintS), 137 (ΔmutS), 149 (CmL05G13), and 143 (GmL07G12). (**h**) Bacterial titers in adult females 35 days after emergence in terms of ntpII gene copies per insect. In (**c**), (**e**) and (**h**), different alphabetical letters indicate statistically significant differences (pairwise Wilcoxon rank sum test: *P* < 0.05).

**Fig. S7.**
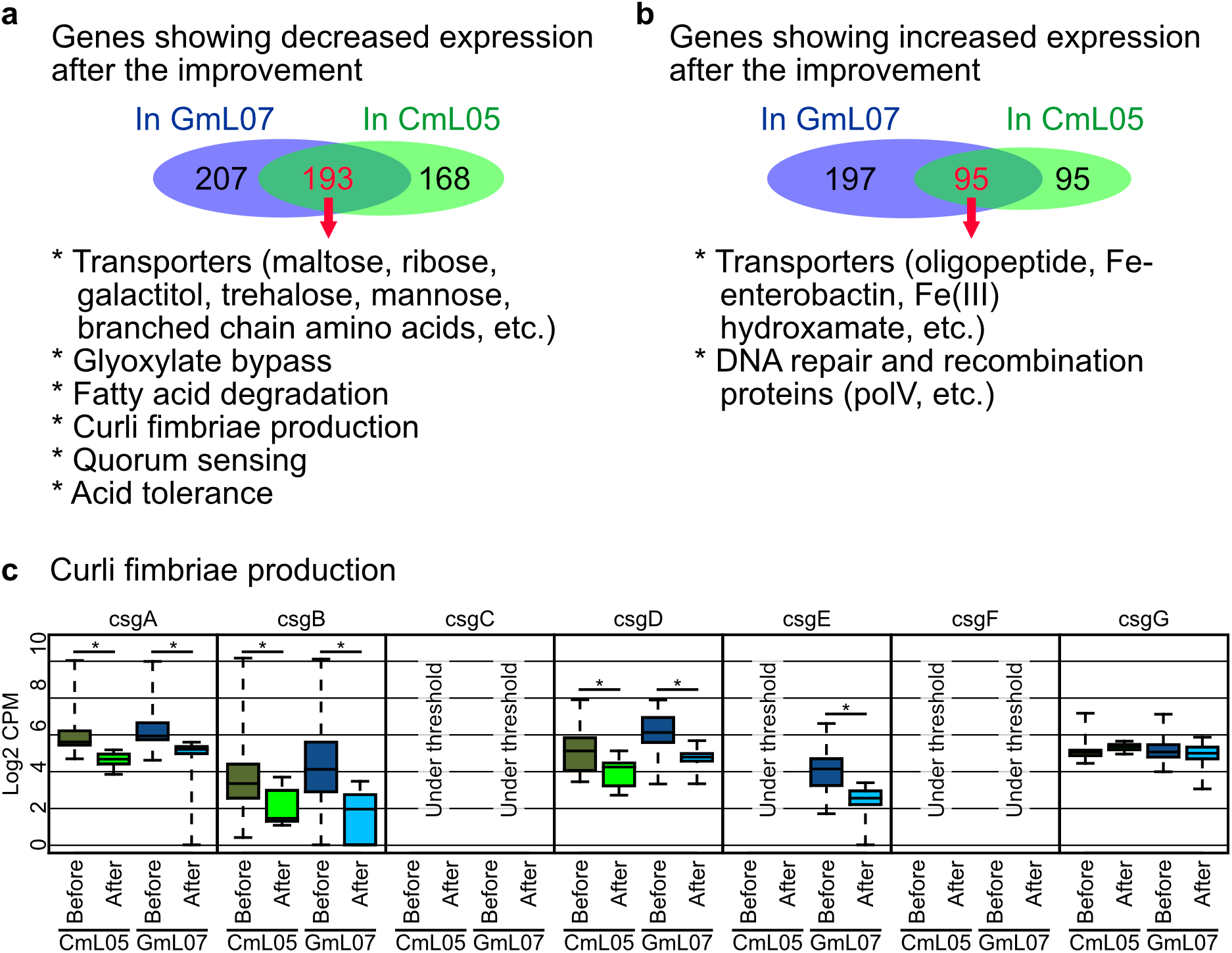
Gene expression changes of evolutionary *E. coli* lines GmL07 and CmL05 before and after improvement of host phenotypes. (**a, b**) Venn diagrams showing down-regulated genes (**a**) and up-regulated genes (**b**) after the improvement of host phenotypes. (**c**) Expression levels of genes involved in extracellular matrix (Curli fimbriae) production before and after the improvement of host phenotypes. Asterisks indicate statistically significant differences (FDR q-value < 0.01).

**Fig. S8.**
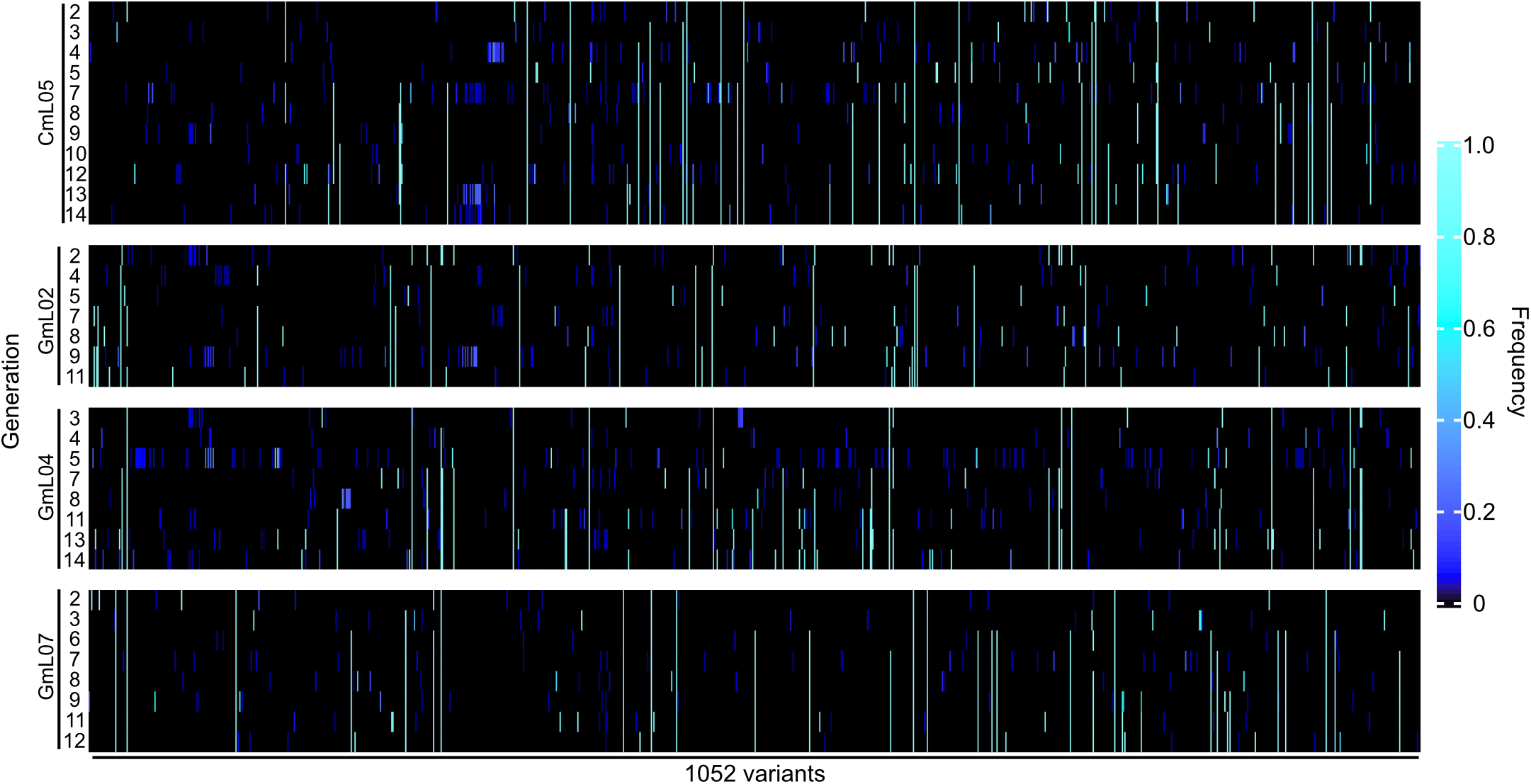
Mutations in the genomes of evolutionary *E. coli* lines CmL05, GmL02, GmL04 and GmL07 in the experimental evolutionary course. Frequencies of 1,052 variants identified in the experimental evolution lines and generations are color-coded. Vertical axis represents the generations of the experimental evolution lines whereas horizontal axis represents an array of 1,052 variants. This figure is the graphical representation of table S7.

**Fig. S9.**
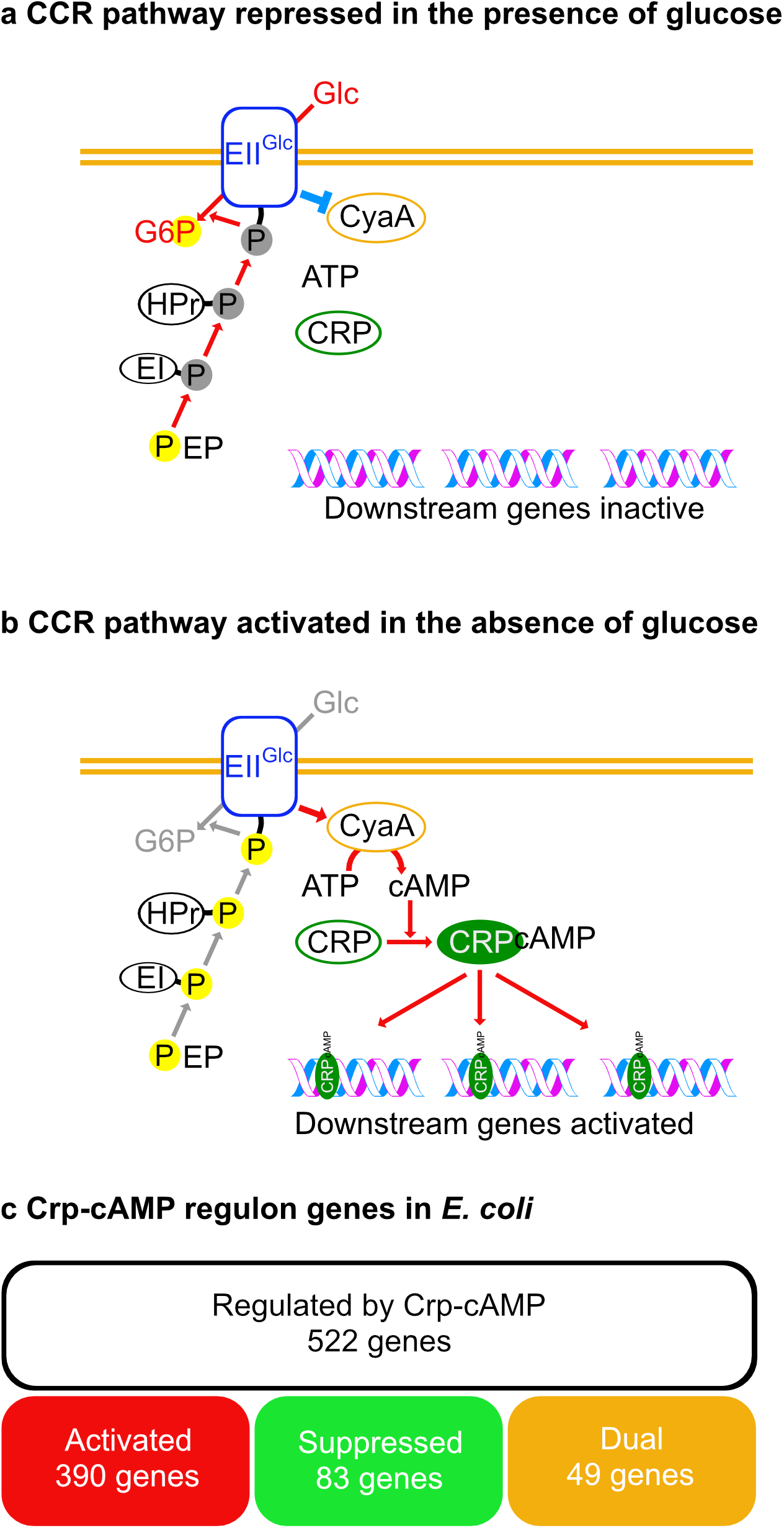
Carbon catabolite repression (CCR) pathway and Crp-cAMP regulon of *E. coli.* (**a**) CCR pathway repressed in the presence of glucose. (**b**) CCR pathway activated in the absence of glucose. (**c**) Number of genes constituting the Crp-cAMP regulon of *E. coli* estimated by RegulonDB (*20*).

**Fig. S10.**
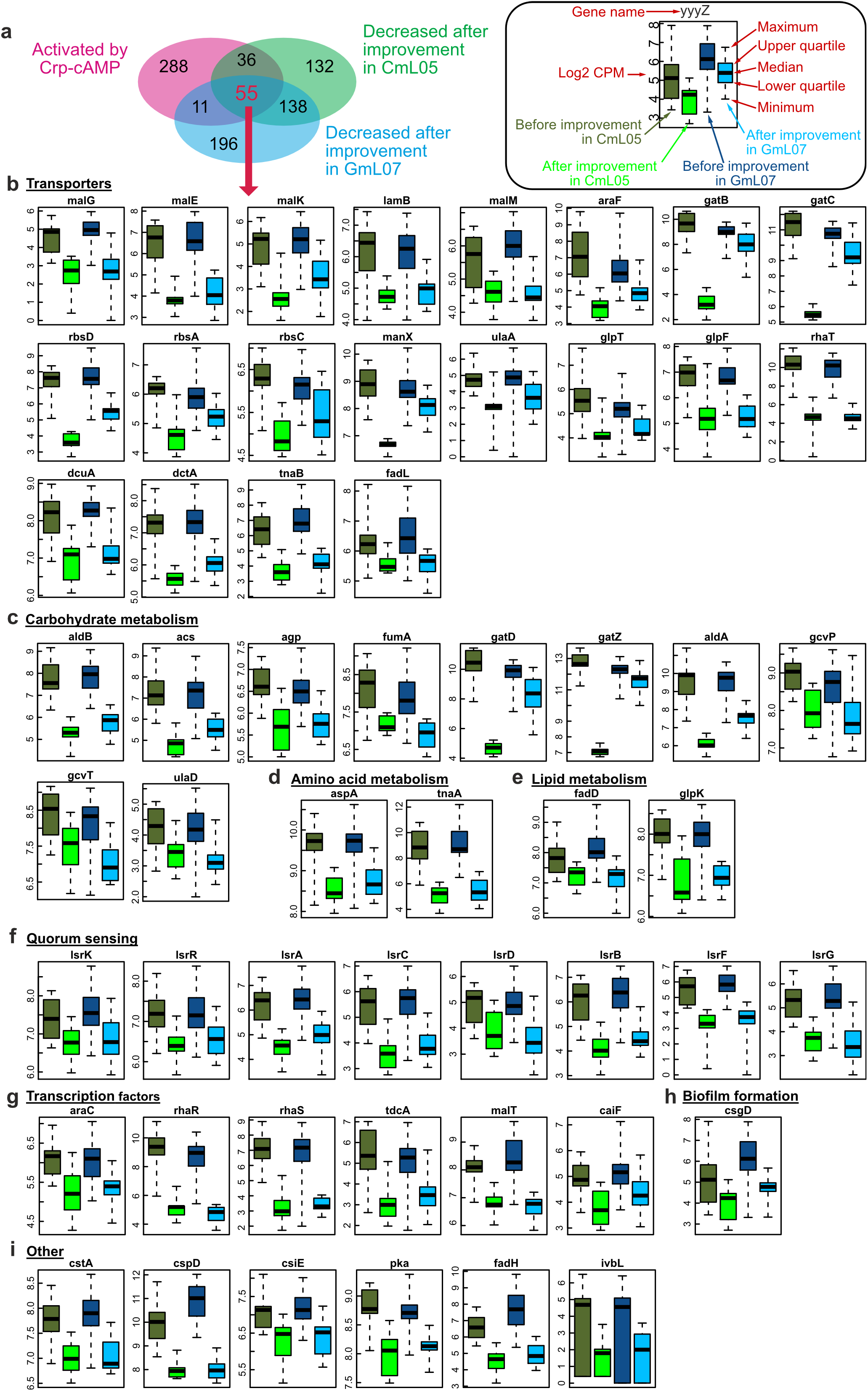
Genes commonly down-regulated in GmL07 and CmL05 after the improvement of host phenotypes, and also down-regulated by disruption of Crp-cAMP in *E. coli*. (**a**) Venn diagram showing the commonly down-regulated genes. (**b-i**) Expression levels of the commonly down-regulated genes in GmL07 and CmL05 after the improvement of host phenotypes. (**b**) Transporter genes. (**c**) Carbohydrate metabolism genes. (**d**) Amino acid metabolism genes. (**e**) Lipid metabolism genes. (**f**) Quorum sensing genes. (**g**) Transcription factor genes. (**h**) Biofilm (= Curli fimbriae) formation genes. (**i**) Other genes.

**Fig. S11.**
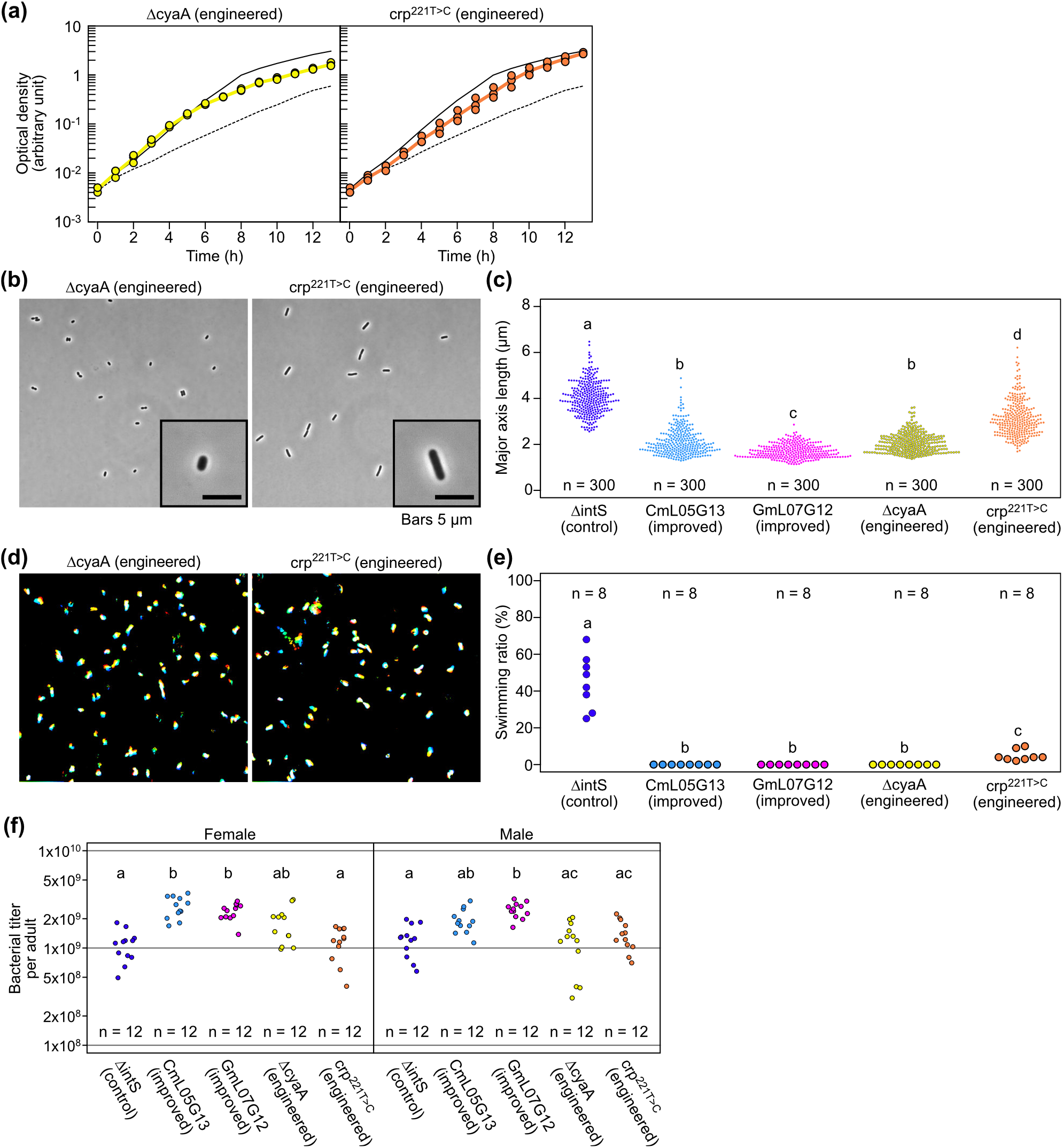
Phenotypic traits of ΔcyaA and crp^221T>C^ mutants of *E. coli.* (**a**) Growth curves (3 replicates each). Upper solid line is the trace of ΔintS growth curve, whereas lower dotted line is the trace of CmL05 growth curve. (**b**) Morphology of bacterial cells. (**c**) Quantification of cell size in terms of major axis length. (**d**) Motility of bacterial cells visualized by rainbow plot for 2 sec. (**e**) Quantification of bacterial motility in terms of number of swimming cells per 100 cells observed. (**f**) Bacterial titers in adult females 35 days after emergence in terms of ntpII gene copies per insect. In (**c**), (**e**) and (**f**), different alphabetical letters indicate statistically significant differences (pairwise Wilcoxon rank sum test: *P* < 0.05).

## Notes

### Competing Interest Statement

The authors have declared no competing interest.

